# An ancient retroviral RNA element hidden in mammalian genomes and its involvement in coopted retroviral gene regulation

**DOI:** 10.1101/2021.03.02.433518

**Authors:** Koichi Kitao, So Nakagawa, Takayuki Miyazawa

## Abstract

Retroviruses utilize multiple unique RNA elements to control several aspects of RNA processing, such as splicing, subcellular export, and translation. However, it is mostly unclear whether such functional RNA elements are present in endogenous retroviruses (ERVs), many of which were inserted into the host genomes millions of years ago. Previously, in human ERV-derived *syncytin-1* gene, we found a *cis*-acting RNA element named SPRE that enhances its protein expression. In this study, we found a 17-nt common sequence in SPRE of *syncytin-1* and another ERV-derived gene, *syncytin-2*, and the sequence is confirmed to be essential for the expression of the proteins. We detected the sequences of SPRE-like elements in 41 ERV families. Though the SPRE-like elements were not found in currently prevailing (*i.e.* exogenous) viral sequences, more than thousands of copies of the elements were found in several mammalian genomes, suggesting the ancient integration and propagation of the SPRE-harboring retroviruses in mammalian lineages. Indeed, other mammalian ERV-derived genes: *mac-syncytin-3* of macaque, *syncytin-Ten1* of tenrec, and *syncytin-Car1* of Carnivora contain the SPRE-like elements, and we validated their function for efficient protein expression by *in vitro* assays. A reporter assay revealed that the enhancement of gene expression by SPRE depended on reporter genes. Moreover, the mutation in SPRE did not affect the gene expression in codon-optimized *syncytin-2*. However, the same mutation in SPRE impaired the gene expression in wild-type *syncytin-2*, suggesting that the SPRE dependency of Syncytin-2 expression is due to the negative factors such as inefficient codon frequency or repressive elements within the coding sequence. These results provide new implications that ERVs harbor unique RNA elements involved in the regulation of ERV-derived genes.

## Introduction

As the traces of ancient organisms remain as fossils, the traces of ancient retroviruses remain as nucleic acid sequences in our host genomes. They are called endogenous retroviruses (ERVs), which are remnants of ancient retroviruses that are incorporated into the genome through infection to host germ cells. ERVs are not mere fossil records. Some of them are still active as host genes or regulatory sequences in the host genome. They have inherited their functions in ancestral viruses: for example, the placental fusogenic Syncytin from the fusogenic envelope protein (1), and lineage-specific host enhancers/promoters from the long terminal repeat (LTR) (2). On the other hand, much is not known about the role of RNA elements in ERVs. Recent studies have shown that several host RNA-binding proteins interact with ERV transcripts (3). However, it is not clear whether unique RNA elements are present in ERVs and what biological significance they have for the hosts.

In order to balance the expression amounts of the viral proteins, RNA elements in retroviruses provide a layer of post-transcriptional regulation. Retroviruses have three major genes, the *gag* gene encoding the major structural protein Gag, *pol* gene encoding polyprotein Pol, consisting of multiple proteins including RNase H, reverse transcriptase and integrase, and *env* gene encoding the envelope protein Env. Some retroviruses have additional genes. uman immunodeficiency virus (HIV)-1 belonging to the Genus *Lentivirus* encodes the regulatory Rev protein that binds to the Rev-responsive element (RRE) in the *env* region (4, 5). Binding of Rev to the RRE facilitates the un-spliced RNA export to the cytoplasm with host factor CRM1/XPO1 (6) as well as the translation of Env and accessory and regulatory proteins (7). Similarly, Rex of human T-lymphotropic leukemia virus 1 belonging to the Genus *Deltaretrovirus* (8) and Rem of murine mammary tumor virus belonging to the Genus *Betaretrovirus* (9) are regulatory proteins that bind to their viral RNAs, allowing efficient viral replication. Mason-Pfizer monkey virus (MPMV) belonging to Genus *Betaretrovirus* does not have regulatory proteins; however, MPMV has an RNA element called the constitutive transport element (CTE). It was initially reported that the CTE has an ability to compensate for Rev-deficient HIV-1 (10). Then, it was revealed that binding of a host protein TAP/NXF1 to the CTE promotes nuclear transport and the translation of un-spliced viral RNA (11, 12). Similarly, NXF1 binding to a viral RNA element called the cytoplasmic accumulation element (CAE) in murine leukemia virus belonging to the Genus *Gammaretrovirus* also promotes the expression of viral proteins (13). Recent comprehensive mutagenesis approaches have revealed that HIV-1 RNA contains many undefined RNA elements required for efficient viral replication (14). Thus, retroviruses have complex RNA elements in their short genomes that allow them to replicate efficiently.

Revealing specific viral RNA elements from ERVs is challenging because accumulated mutations have erased functional RNA elements. Exceptionally in HERV-K families, which are young ERVs retaining intact viral ORFs and shows polymorphic loci in human genomes (15, 16), the post-transcriptional roles of their RNA-binding regulatory protein, Rec and its binding RNA element have been demonstrated (17). For investigating more ancient viral RNA elements, co-opted viral genes could provide important clues because viral RNA elements have been possibly conserved with their ORFs to regulate the gene expression. *Syncytin-1* is an *env* gene of ERVWE1 and contributes to the cell fusion for the differentiation of multinucleated syncytiotrophoblast in human placenta (18, 19). We have previously reported that an RNA element located in the 3′ end of the ORF and 3′ untranslated region (3′ UTR) of human *syncytin-1* is important for its protein expression, and was named *<u>s</u>yncytin* <u>p</u>osttranscriptional regulatory <u>e</u>lement (SPRE) (20). Indeed, human *syncytin-2*, another *syncytin* gene derived from an *env* gene of ERVFRDE1 (21) also appears to contain functional motifs in their 3′ UTR necessary for efficient expression (20). Such RNA elements would enable us to access to the RNA regulatory mechanisms of ancient retroviruses.

In this study, a hidden Markov model (HMM)-based sequence search in ERV database revealed the core motifs of SPRE. We found that the defined SPRE-like elements were widespread in distinct ERV families but not in extant viruses. The SPRE-like elements were also detected in three non-human *syncytin* genes. A reporter assay verified their functionality and elucidated the unique features that nucleotide sequences of the target genes affect the SPRE activity. These results provide new insights into ancient retroviral post-transcriptional regulation as well as its involvement to the co-opted genes from ERVs.

## Results

### The SPRE-core motif is functionally essential for the SPRE activity

A partial sequence of 3′ end of the ORF and the subsequent 3′ UTR of human *syncytin-1* (68-nt) and a partial sequence of 3′ UTR of *syncytin-2* (400-nt) increased protein expression when inserted into the 3′ UTR of an HIV-1 Gag expression plasmid (20) (Fig 1A). We speculated that these sequences share functional RNA motifs. To explore the essential motif(s), the two regulatory sequences were aligned and compared. We revealed that a 17-nt common sequence (5′-TCAGCAGGAAGCAGTTA-3′) is shared between *syncytin-1* and *syncytin-2* (Fig 1B). Next, we examined whether the common sequence is essential for the expression of Syncytin-1 and Syncytin-2. We generated expression plasmids by cloning *syncytin-1* and *syncytin-2* with their 3′ UTRs and introduced several mutations into the 17-nt common sequence (Fig 1C). Since this sequence overlaps with the *syncytin-1* ORF, we generated mutants that avoid any amino acid substitutions. In this experiment, we did not add a peptide tag at their C-termini to detect protein expression because it may alter the SPRE-activities. Therefore, their protein expression levels were evaluated by a cell fusion-dependent luciferase assay utilizing the property that both Syncytin-1 and Syncytin-2 induce strong cell-fusion. As a result, mutations in the common sequence markedly reduced cell-fusion activities for both Syncytin-1 and Syncytin-2 (Fig 1D), suggesting the importance of the common sequence for the SPRE activity.

**Fig 1.**
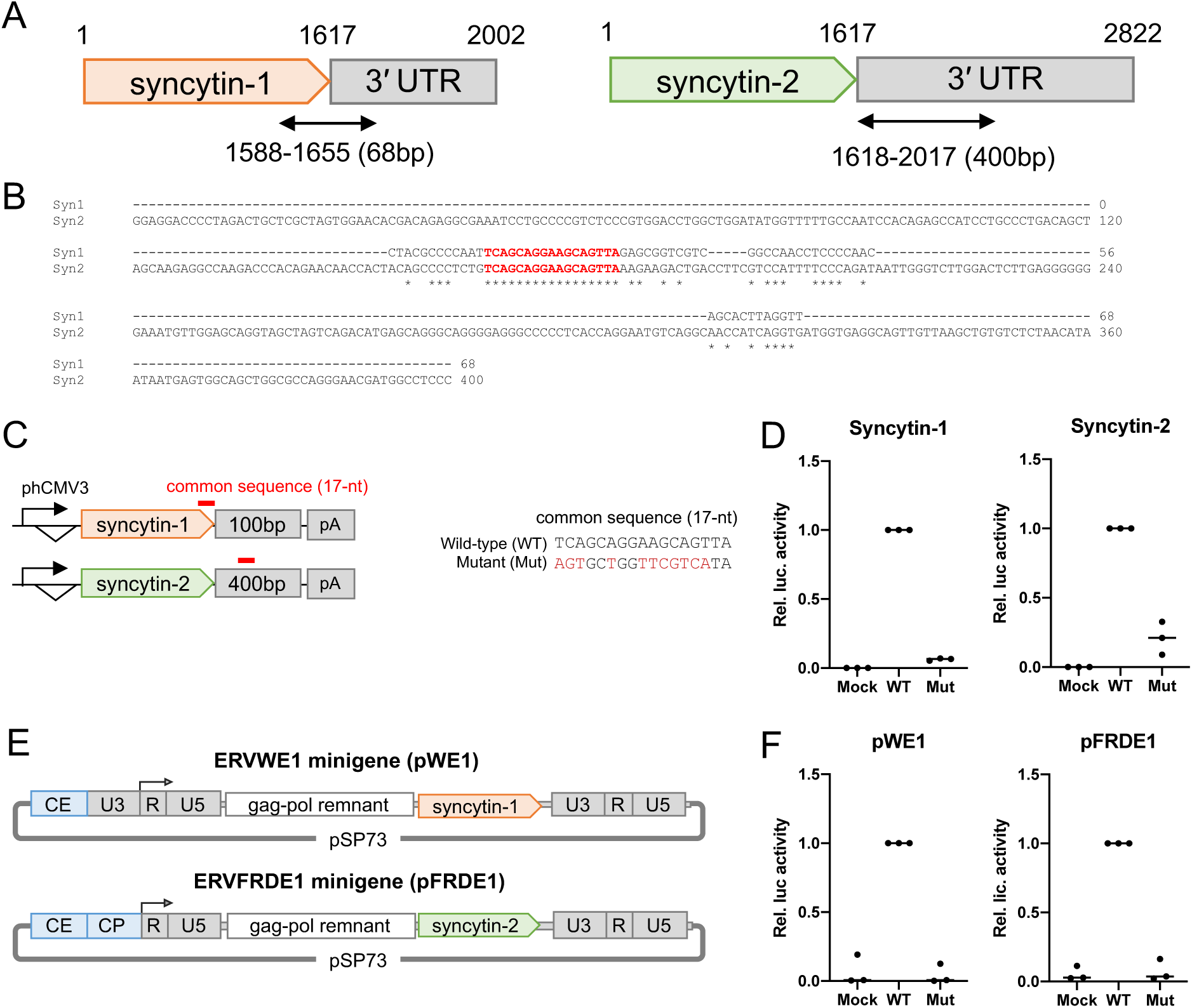
Identification of the 17-nt SPRE-core motif in *syncytin-1* and *syncytin-2*. (A) Location of the SPRE sequences which increase protein expression of HIV-1 Gag in human *syncytin-1* and *syncytin-2* (68-nt and 400-nt, respectively). (B) Alignment of the two sequences. The 17-nt common sequence is represented by the square. (C) Schematic representation of the Syncytin-1 and Syncytin-2 expression plasmids and the sequence of the SPRE mutants. (D) Fusion-dependent luciferase assay in 293T cells transfected with Syncytin-1 and Syncytin-2 expression plasmids. WT; wiled-type sequence, Mut; SPRE mutant. Each value was normalized by the luminous intensity of WT. Individual data were indicated as points, and medians were indicated as bars. (E) Schematic representation of minigene plasmids of *syncytin-1*; pWE1 and *syncytin-2*; pFRDE1. The CMV enhancer and promoter were introduced their upstream to enhance the gene expression. CE; CMV enhancer, CP; CMV promoter. (F) Fusion-dependent luciferase assay of 293T cells transfected with pWE1 and pFRDE1.

To infer the importance of the common sequence in native mRNA forms, which may include various intrinsic *cis*-acting elements affecting the SPRE activity, we generated proviral minigene plasmids of *syncytin-1* and *syncytin-2,* including their 5′ and 3′ LTRs (Fig 1E). Then, we introduced mutations into the common sequence and compared their protein expression levels by performing the cell-fusion-dependent luciferase assay. As a result, we found that mutations into the common sequence impaired the cell-fusion activity (Fig 1F). Based on these results, we named the common sequence (5′-TCAGCAGGAAGCAGTTA-3′) as the SPRE-core motif that is functionally essential for the SPRE activity.

### SPRE-like elements in ERV families

To investigate whether the other ERVs also contain SPREs, we searched for the SPRE-core motif against the Dfam database version 3.2, an open collection of transposable elements (22). The 17-nt of the SPRE-core motif is too short to avoid non-specific hits with statistical significance. To reduce the non-specific hits, we performed a two-step search strategy (Fig 2A). In the first step, we focused on the Dfam entries classified as ERVs, consisting of three groups: 447 families of ERV1 (*Gamma-* and *Epsilonretrovirus*-like), 379 families of ERV2 (*Alpha-*, *Beta-*, and *Deltaretrivirus*-like), and 270 families of ERV3 (HERV-L-like). Furthermore, we calculated relative positions of the hits to select ones near their 3′-termini because SPREs thought to be located in 3′ UTRs of *env* genes. To determine the relative positions, each hit position to the SPRE-core motif was normalized by the nucleotide length of each ERV and was scored from 0 to 100 (from 5′ to 3′ ends, respectively). As a result, the hit positions were highly biased towards the 3′-terminal regions, particularly for ERV1 group (Fig 2B and S1 Table). We considered that the hits satisfying the following conditions were genuine SPREs: 1) belonging to the ERV1 group and 2) relative position scores were more than 97. As a result, we regarded the 34 families satisfying the above conditions as SPRE-harboring ERVs. In the second step search, we collected core motifs and their flanking sequences of the 34 ERVs from the first step search and aligned them (S1 Fig). Then, we constructed a hidden Markov model (HMM) profile based on the alignment and searched for the element in 6,915 families in Dfam including ERVs and non-ERV transposons using the nhmmer program in the HMMER (23) (Fig 2A). By this search, we obtained 41 SPRE-harboring ERV families; 34 hits belonged to the ERV1 group, which were the same as the result of the first step search, and 7 hits belonging to the ERV2 group (Fig 2C). The SPRE-like elements were not detected in non-ERV transposable elements. The sequence logo of the alignment showed two C-rich regions in both 5′- and 3′-flanking regions of the SPRE-core motif (Fig 2C), indicating that SPRE consists of three motifs; the SPRE-core motif and two C-rich regions.

**Fig 2.**
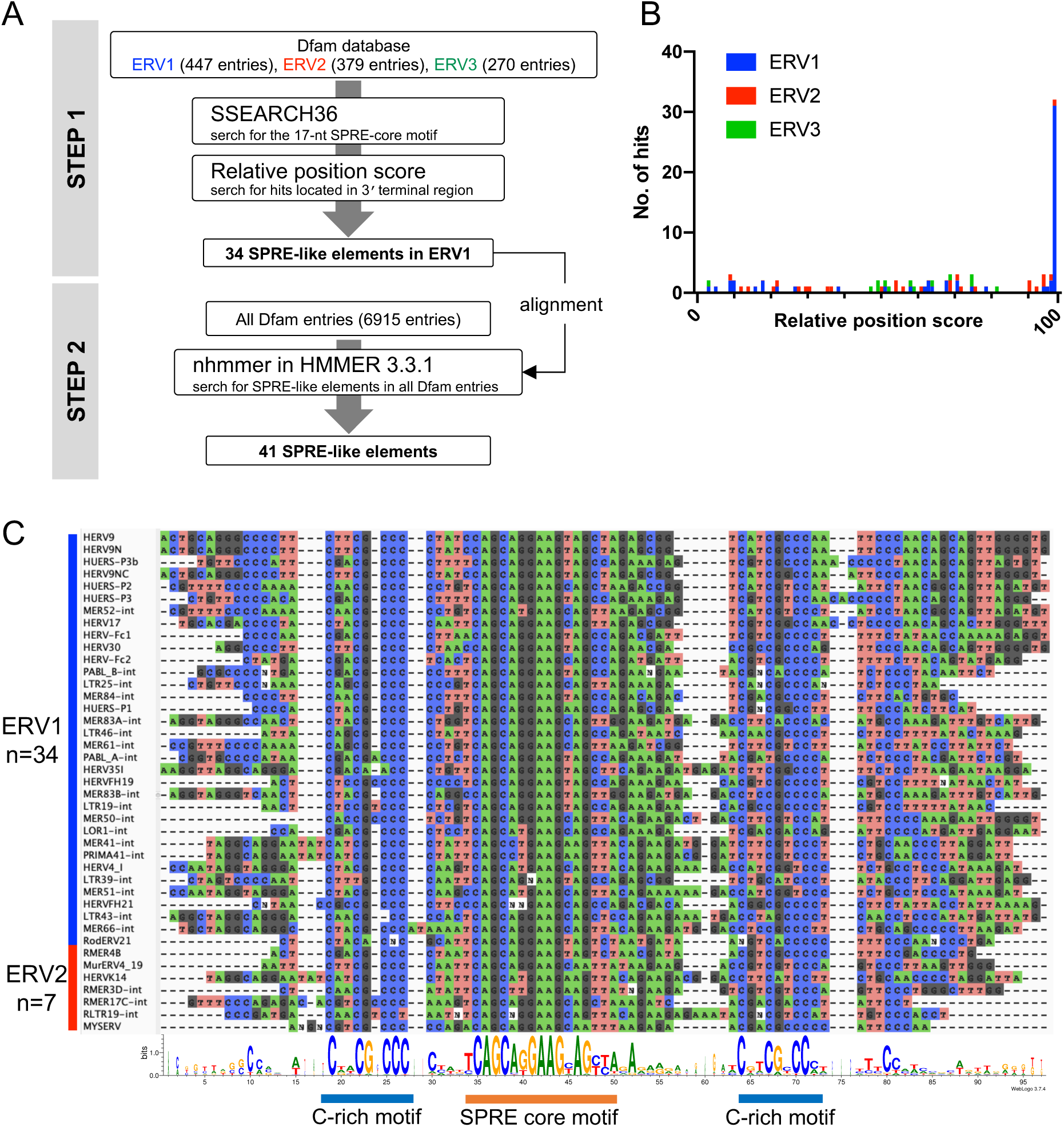
The sequence search for the SPRE-like elements in ERV families. (A) Schematic representation of the two-step search for SPRE in Dfam database. (B) Relative position of the hits to the SPRE-core motif in ERV consensus sequences. Score 0 represents 5′-terminus and score 100 represents 3′-terminus of each ERV sequence. (C) The Sequence alignment and the sequence logo of 41 SPRE-like elements. The SPRE-core motif and C-rich motifs were indicated.

### No exogenous viruses with the SPRE-like elements and a number of the elements in host genomes

To investigate whether currently prevailing (*i.e*. exogenous) retroviral genomes contain SPRE-like elements, we first searched for viral sequences obtained from all viral nucleotide sequences deposited in the NCBI virus database (https://www.ncbi.nlm.nih.gov/labs/virus/) using the HMM profile constructed based on the alignment of 41 SPRE-like elements shown in Fig 2C. We found that there were no exogenous viruses that contain SPRE-like elements. We only obtained a hit with a threshold (Accession Number: AF127229.1, E-value = 9.3-e7); however, it was multiple sclerosis retrovirus, derived from ERV-W-related sequences (24). Next, we searched for the SPRE-like elements in genomes of 318 species of mammals and 499 species of birds available in the NCBI genome database (https://www.ncbi.nlm.nih.gov/genome/). The SPRE-like elements were found in almost all mammalian genomes examined, except for *Erinaceus europaeus* (European hedgehog). The number of hits varied greatly among species within the same orders. For example, in *Rodentia*, we identified up to 9691 hits in the genome of *Microtus arvalis*, while only two hits were detected in the genome of *Sigmodon hispidus* (Fig 3, S2 Table). This suggests that the copy numbers of the SPRE-like elements have been increasing rapidly and independently in many mammalian lineages. In contrast, in the avian genomes, only six species were found to have the SPRE-like elements (Fig 3). It should be noted that we could not rule out the possibility of contamination of human or mouse DNA for the following reasons: all of 14 hits with flanking sequences found in avian genomes showed high similarity to sequences of *Homo sapiens* or *Mus musculus,* and seven hits were located within short contigs (<5000 bp) within each genome. These results indicate that while the SPRE-like elements are not present in the currently exogenous retroviruses, infections and/or transpositions of the SPRE-harboring retroviruses have been repeated through mammalian evolution.

**Fig 3.**
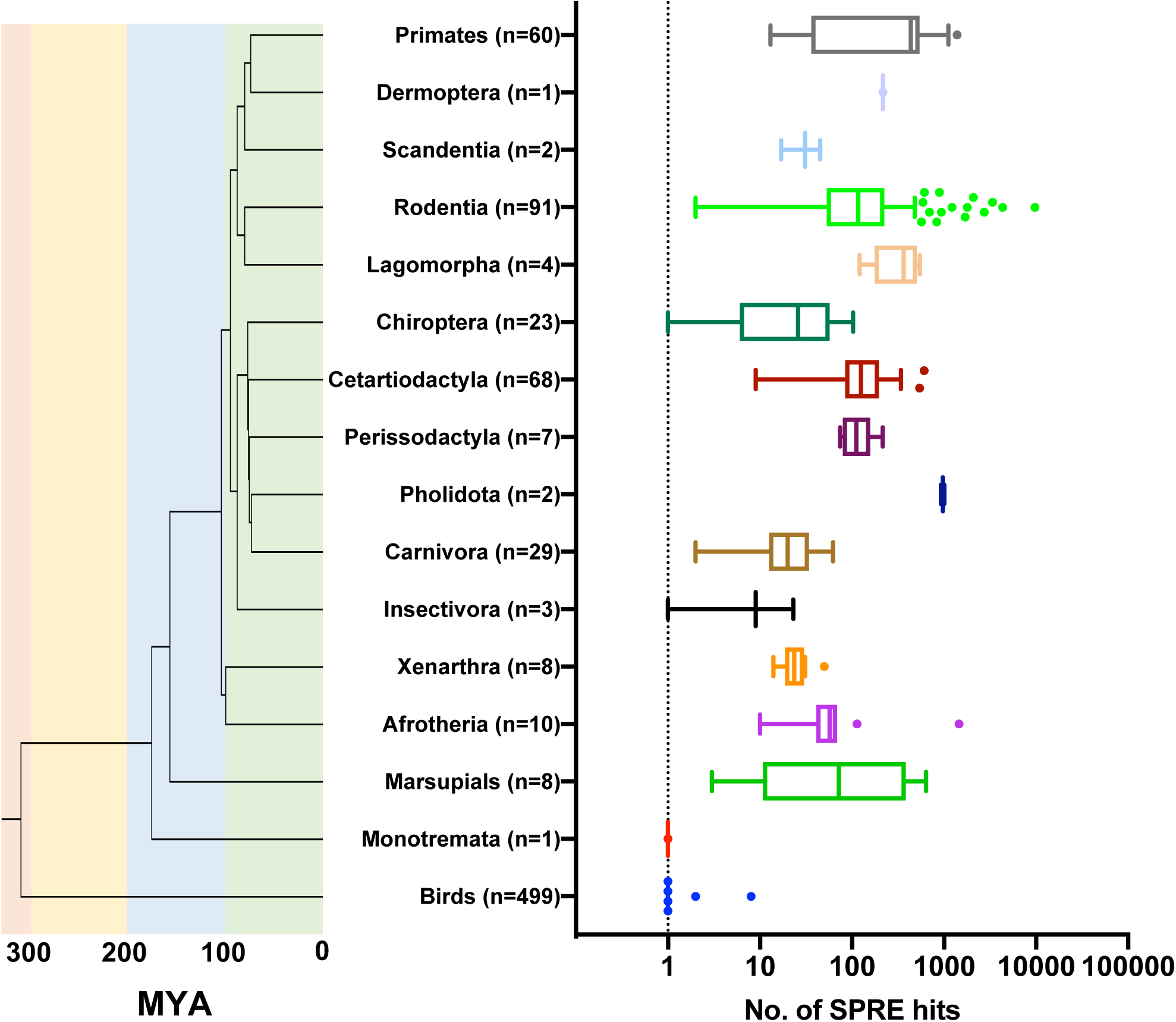
Numbers of SPRE-like elements in host genomes. Number in each lineage indicates species used for the genomic search of SPRE.

### Various mammalian *syncytin* genes retain the SPRE-like elements

Next, we attempted to determine whether the SPRE-like elements are involved in the co-option of ERV-derived genes other than *syncytin-1* and *syncytin-2*. For this purpose, we examined other mammalian *syncytin* genes, which were acquired from various distinct ERVs in an independent manner (25, 26). We conducted a HMM search for the SPRE-like elements in ORFs and 1000-nt of 3′ flanking sequences of *mac-syncytin-3* in macaque (27), *syncytin-A*, and *-B* in mouse (28), *syncytin-Mar1* in squirrel (29), *syncytin-Ory1* in rabbit (30), *syncytin-Rum1* (31) and *fematrin-1* in cow (32), *syncytin-Car1* in dog (33), *syncytin-Ten1* in tenrec (34), and *syncytin-Opo1* in opossum (35). As a result, we found the SPRE-like elements in the 3′ UTRs of *mac-syncytin-3*, *syncytin-Ten1*, and *syncytin-Car1* (Fig 4A). In *mac-syncytin-3*, cell fusion activity was observed with the addition of the 3′ UTR of *mac-syncytin-3*, but no cell-fusion activity was observed without the 3′ UTR or with a mutation in the SPRE-core motif (Fig 4B). We also evaluated the function of 3′ UTR in the *syncytin* genes where the SPRE-like elements are not found using *syncytin-A* in mouse. It was found that Syncytin-A caused cell fusion irrespective of the presence of 3′ UTR, suggesting that *syncytin-A* does not have crucial RNA elements in the 3′ UTR (Fig 4C). The *syncytin* genes containing the SPRE-like elements are not phylogenetically related as illustrated by a phylogenetic tree (Fig 4D).

**Fig 4.**
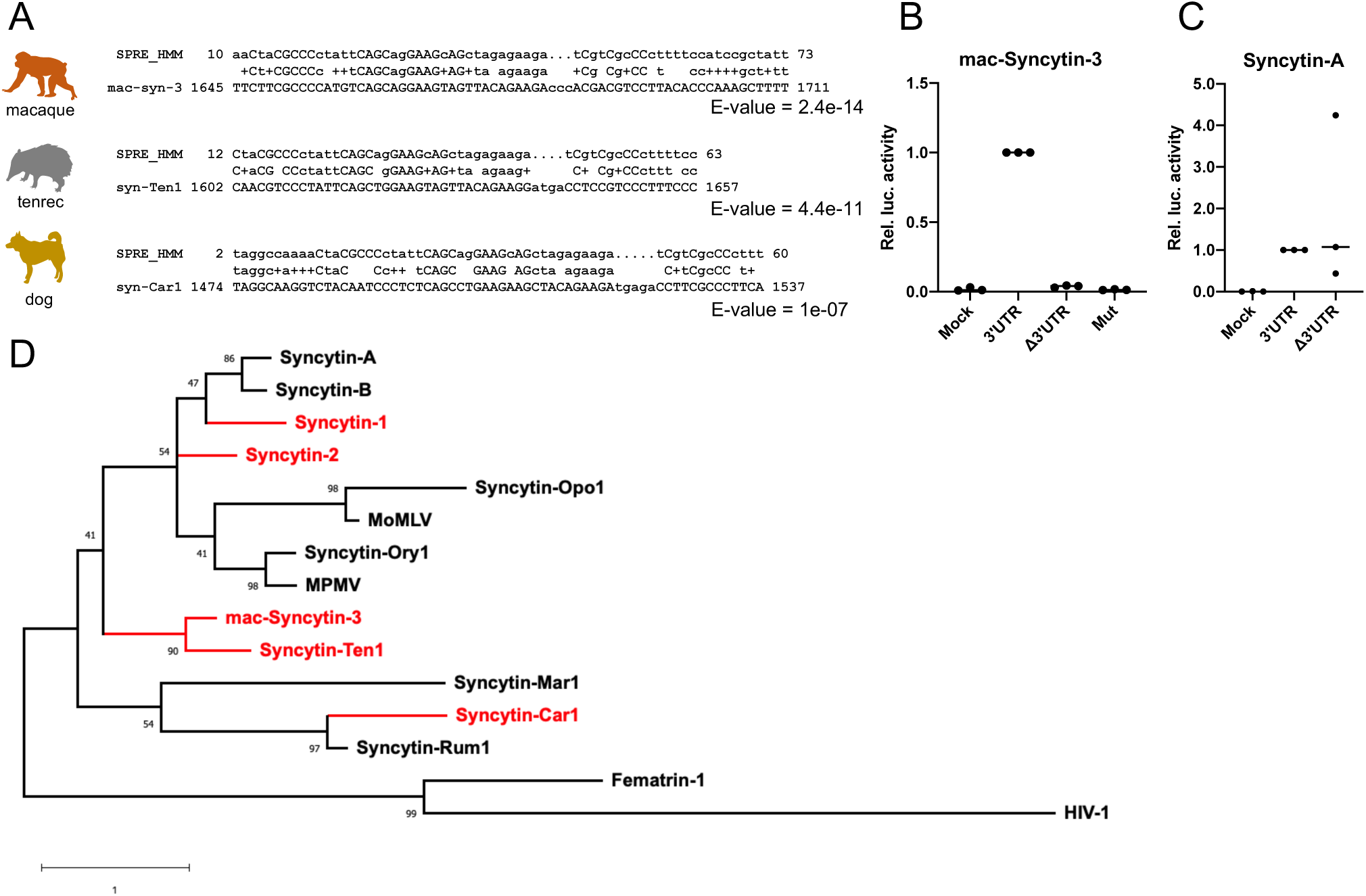
Identification of the SPRE-like elements in different mammalian *syncytin* genes. (A) Alignment of the HMM profile and the SPRE-like elements in *mac-syncytin-3*, *syncytin-Ten1*, and *syncytin-Car1*. Numbers in *syncytin* genes indicate the nucleotide position from the first nucleotide of the start codon. (B and C) Fusion-dependent luciferase assay in mac-Syncytin-3 (B) and Syncytin-A (C) using 293T cells. Each *syncytin* gene was cloned into phCMV3 plasmid with or without 3′ UTR. The SPRE mutation similar to Fig 2A was introduced into *mac-syncytin-3*. (D) Maximum likelihood phylogenetic tree of TM domains in Syncytin proteins and other retroviral Env proteins. The bootstrap support values are displayed at the nodes. SPRE-harboring Syncytins are indicated as red color. Accession numbers of sequences are listed in Table S3.

### Functional analysis of the SPRE-like elements by the reporter assay

Next, we tried to verify the functional activities of these SPRE-like elements using a reporter assay with HIV-1 Gag as a reporter protein. Since SPRE of *syncytin-1* (SPRE-syn1) was identified as a 68-nt sequence including the SPRE-core motif (17-nt) with 5′-flanking (12-nt) and 3′-flanking (39-nt) sequences for enhancing HIV-1 Gag expression (20), we also constructed a reporter plasmid containing SPRE of *syncytin-2* (SPRE-syn2) in the same manner (Fig 5A). We applied a recently developed luciferase system called HiBiT (Promega) to quantify the protein amounts by measuring the luminous activities, and HIV-1 Gag was fused with the C-terminal HiBiT tag (HG-HiBiT). We then inserted the SPRE-syn1 and SPRE-syn2 into the 3′ UTR of the HG-HiBiT. As expected, the expression of HG-HiBiT was increased by the insertion of two SPREs, whereas the mutation into the SPRE-core motif abolished the effects (Fig 5B). We also found that the SPRE-core motif alone did not increase the protein expression level (Fig 5B). To verify the importance of C-rich motifs shown in Fig 2C, we constructed the C-to-G mutants of the C-rich motifs in SPRE-syn1 and SPRE-syn2 (Fig 5A). These mutations impaired the enhancement of gene expression in both SPRE-syn1 and SPRE-syn2, suggesting that the C-rich region is crucial in the SPRE activity (Fig 5C and D). To test the functional activities of the SPRE-like elements found in other mammalian *syncytin* genes (Fig 4A), the SPRE-like elements from *mac-syncytin-3*, *syncytin-Ten1*, and *syncytin-Car1* were inserted into HG-HiBiT (Fig 5A). We found that all three SPRE-like elements enhanced the protein expression of the HIV-1 *gag* construct (Fig 5E).

**Fig 5.**
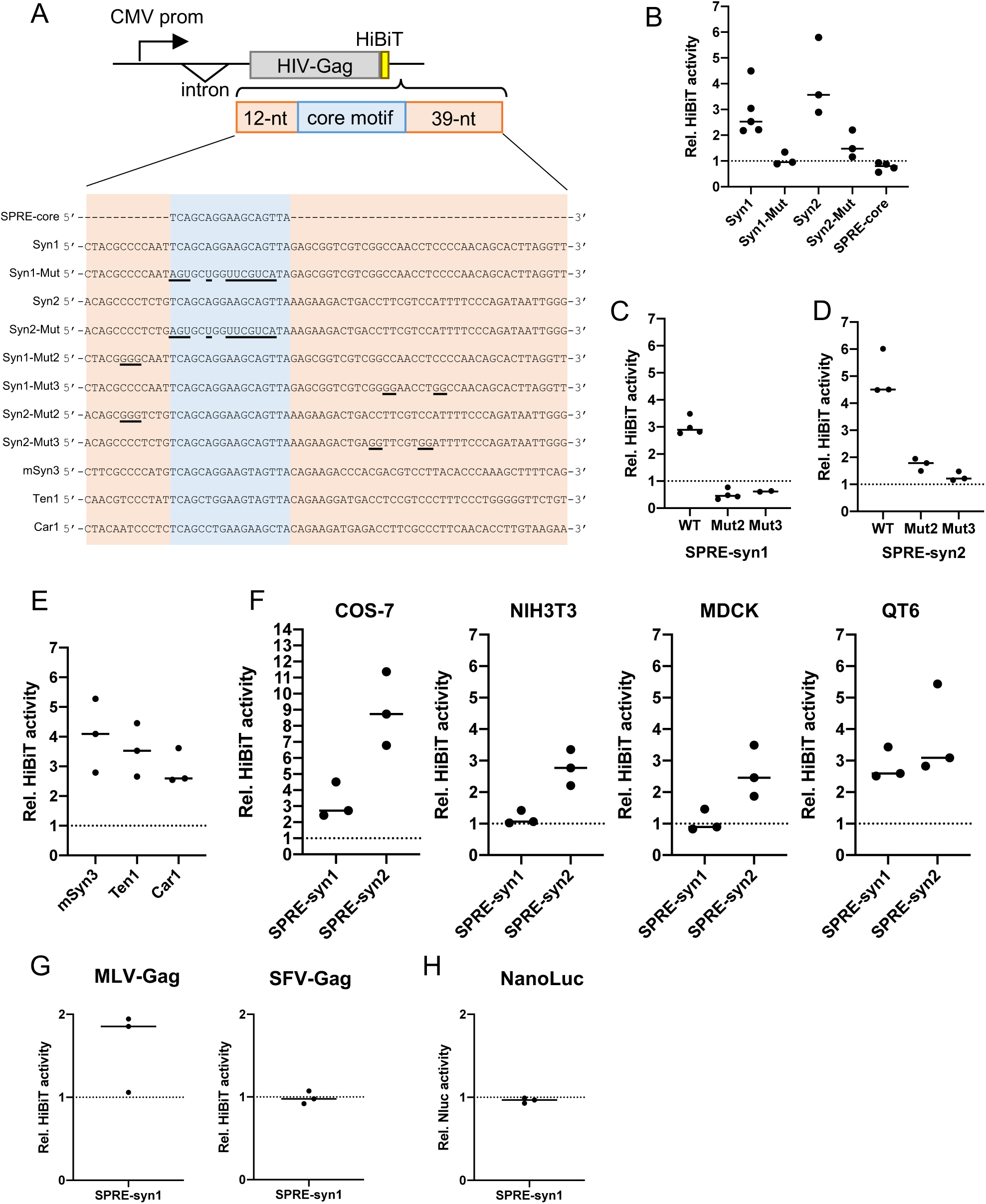
Functional analysis of SPRE-like elements using a reporter assay. (A) Schematic representation of HIV-1 Gag-HiBiT (HG-HiBiT) reporter plasmid. Introduced mutations were indicated by underlines. (B) Luminous intensities were detected in 293T cells transfected with HG-HiBiT reporter plasmids. Each value was normalized by the luminous intensity from HG-HiBiT expression plasmid without an additional sequence in 3′ UTR. (C and D) Functional investigation of C-to-G mutants of SPRE-syn1 (C) and SPRE-syn2 (D). (E) Functional investigation of the SPRE-like elements of *mac-syncytin-3*; mSyn3, *syncytin-Ten1*; Ten1, and *syncytin-Car1*; Car1 by the HG-HiBiT reporter assay in 293T cells. (F) Functional investigation of SPRE-syn1 and SPRE-syn2 in COS7 (African green monkey), NIH3T3 (mouse), MDCK (dog), and QT6 (quail) by the HG-HiBiT reporter assay. Each value was normalized by the luminous intensity from HG-HiBiT expression plasmid without an additional sequence in 3′ UTR in each cell line. (G) Functional investigation of the SPRE activities in MLV-Gag and SFV-Gag. Each value was normalized by the luminous intensity of respective reporter without an additional sequence in 3′ UTR. (H) Functional investigation of the SPRE activities in NanoLuc.

Next, we tested the SPRE activity in mammalian and avian cell lines other than 293T. We conducted HG-HiBiT reporter assay in African green monkey (COS-7), mouse (NIH3T3), dog (MDCK), and quail (QT6) cell lines. Though SPRE-syn1 did not increase the HG-HiBiT expression in NIH3T3 and MDCK, SPRE-syn2 was functionally active in all cell lines examined in this study (Fig 5F). These results indicate that SPRE is functional in a wide range of host species.

Considering the fact that the SPRE-like elements were found in various ERVs and *syncytin* genes, they may enhance a wide variety of reporter genes not limited to HIV-1 Gag. To verify this hypothesis, SPRE-syn1 was inserted into the 3′ UTR of Gag proteins of murine leukemia virus (MLV) and simian foamy virus (SFV) with HiBiT, and we measured their protein expression levels by HiBiT luciferase activity. Unexpectedly, the levels of gene expression enhancement by SPRE-syn1 were different among reporter genes; weak enhancement in MLV and no effect in SFV (Fig 5G). SPRE-syn1 was also added downstream of NanoLuc as a non-viral gene, but SPRE-syn1 did not affect the NanoLuc expression (Fig 5H). These data suggest that SPRE activity is dependent on the reporter genes.

### The SPRE activity depends on the ORF sequences

We hypothesized that the SPRE activity depends on the nucleotide sequences of the ORF, not on the reporter proteins. To test this hypothesis, we modified the nucleotide sequence of *syncytin-2* ORF without any amino acid changes by codon optimization in the minigene described in Fig 1E as follows: the wild-type (WT), codon-optimized (CO), and two chimeric sequences of WT and CO (Chimera-A and -B) (Fig 6A). *Syncytin-1* was not used in this analysis because its SPRE is located in its ORF. Cell-fusion dependent luciferase assays revealed that the gene expression of CO-*syncytin-2* was not affected by the mutation in the SPRE-core motif. In the two chimeras, their expression levels were decreased by the SPRE mutation, but the effects were smaller compared to WT-*syncytin-2* (Fig 6B). These data suggest that the SPRE supports the efficient expression of ERV-derived genes, whose codon usages were not optimized for humans.

**Fig 6.**
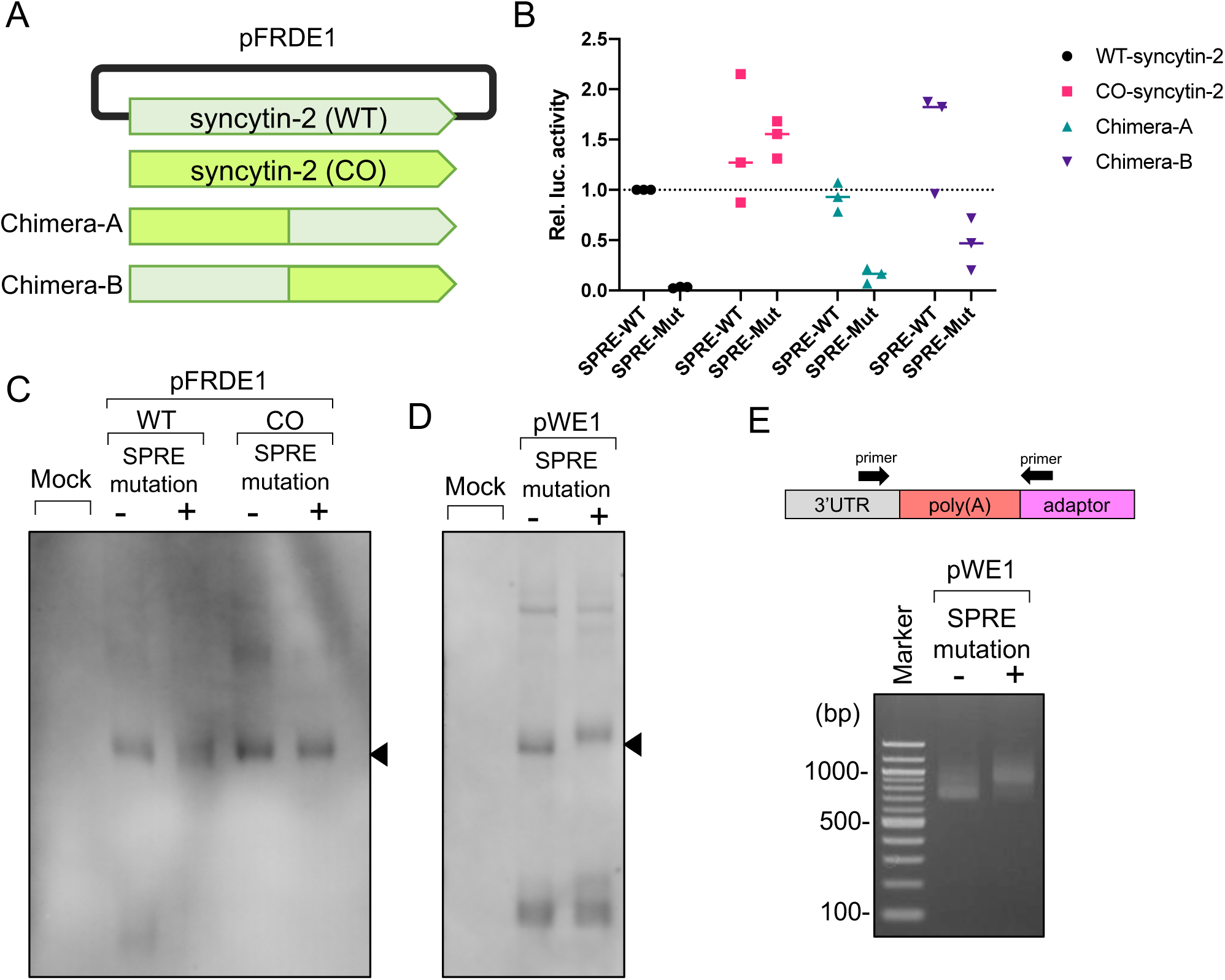
Dependency on the ORF sequences in the SPRE activities. (A) Schematic representation of codon-optimized (CO) *syncytin-2* and chimera *syncytin-2* expression plasmids. The wild-type (WT) *syncytin-2* in pFRDE1 indicated in Fig 1E was replaced by the CO-syncyint-2. (B) Fusion-dependent luciferase assay in WT-*syncytin-2*, CO-*syncytin-2*, Chimera-A, and Chimera-B, with wild-type SPRE (SPRE-WT) or mutated SPRE same as Fig 1C (SPRE-Mut). Each value was normalized by WT-*syncytin-2* with SPRE-WT, and individual data points were indicated as points, and medians were indicated as bars. (C) Northern blot analysis in transcripts from 293T cells transfected with pFRDE1 minigenes; WT-*syncytin-2* and CO-*syncytin-2*, with or without the SPRE mutation. Block arrow indicates the *syncytin-2* mRNA. (D) Northern blot analysis of pWE1 with or without the SPRE-core motif mutation. Block arrow indicates the *syncytin-1* mRNA. (E) Poly(A) tail-targeted PCR in pWE1 transcripts.

To examine the effects of SPRE mutations on the mRNA amounts and sizes, we performed Northern blot analysis using 293T cells transfected with *syncytin-2* minigenes. As a result, the amounts of transcripts slightly decreased by the SPRE mutation in WT-*syncyin-2*, and no significant changes in the size of transcripts were observed (Fig 6C). In *syncytin-1*, the amounts of transcripts were also slightly decreased, but the transcript’s size was increased by the SPRE mutation (Fig 6D). We hypothesized that this band-shift was due to the longer length of poly(A) tail. We conducted a poly(A) tail-targeted PCR and revealed that the SPRE mutant showed longer poly(A) tail length in *syncytin-1* mRNA (Fig 6E). Several studies have reported that highly expressed and well-translated transcripts have short poly(A) tails, probably because the poly(A) tails are decomposed in combination with the active translation and are bound by a minimal number of poly(A) binding proteins (36). Therefore, the longer poly(A) tail produced by the SPRE mutation may suggest a reduced translation efficiency of *syncytin-1* mRNA.

## Discussion

In this study, we characterized a functional retroviral RNA element, termed SPRE, that is found in distinct ERV families, including the ERV-derived *syncytin* genes. In particular, a 17-nt (5′-TCAGCAGGAAGCAGTTA-3′) was identified as the functional core motif of this element (Fig 1). The discovery of the SPRE-core motif enabled the sequence-based searches and revealed that 41 ERV families harbor the SPRE-like elements (Fig 2). The alignment of the SPRE-like elements revealed that SPRE consists of the SPRE-core motif and its upstream and downstream of two C-rich motifs (Fig 2C). Given that the mutagenesis for each motif results in loss of SPRE activity (Fig 5A-D), it is likely that the SPRE activity requires multiple proteins that bind to each motif and formation of a “SPRE-proteins complex”. The RNA secondary structures are also generally crucial for the function of RNA elements. However, our prediction of RNA secondary structures of the SPRE-like elements in *syncytin* genes varied greatly (S1 Fig). The specific secondary structures may not contribute to the function of SPRE, or common structures may be formed by protein binding *in vivo* that cannot be revealed by the predictions based on the nucleotide sequences alone.

We found the SPRE-like elements in the genomes of a variety of mammalian species (Fig 3). While the SPRE-like elements were detected in all mammalian orders, their copy numbers varied greatly among species even within the same orders. Because the SPRE-like elements were not found in non-ERV transposons (Fig 2), the increasing copy numbers of SPRE are mainly due to retrotransposition of ERVs and/or horizontal transmission by exogenous retroviral infection. Considering the repeated invasion into the genomes and the resultant burst of their copy numbers in a wide variety of mammalian lineages, the ancient SPRE-harboring retroviruses may have been prospering viral group(s). It should be noted that the “SPRE-harboring retroviruses” do not indicate a single viral clade, as the SPRE-like elements were identified not only from the ERV1 group but also from the ERV2 group (Fig 2C). Previous attempts of systematic classification of human ERVs revealed their mosaic structures between families and even distinct classes due to recombination (37). Therefore, it is likely that the SPRE-like elements were also inherited from one retrovirus to another retrovirus through recombination. Considering the fact that the SPRE-like elements were found in the genomes of hundreds of mammalian species, there could be many unknown ERVs containing SPRE-like elements in non-model mammalian genomes as well. As more mammalian ERVs are profiled, the evolution of the SPRE-harboring retroviruses will be revealed in more detail.

Despite the ancient prosperity of the SPRE-harboring retroviruses, SPRE has not been identified among the current prevailing exogenous retroviruses. Considering the fact that SPRE is functional in mammalian and avian cells (Fig 5F), SPRE has the potential to give a functional advantage to exogenous retroviruses infecting a wide variety of host species. Therefore, the reason why SPRE is not present in the currently exogenous retroviruses is a puzzling question. One possibility is that the host-virus arms races between hosts and ancient retroviruses drove dynamic exchanges of RNA regulatory mechanisms in retroviruses, rendering SPREs obsolete. An example of conflicts between hosts and ancient retroviruses are have previously been presented in the evolution of a member of host antiviral genes, *APOBEC3*, in which increased repertoire of *APOBEC3* genes is positively correlated with the expansion of ERVs (38). Though whether there are some host defense mechanisms against SPRE is unknown at present, the evolutionary arms race could explain the complete disappearance of SPRE from prevailing exogenous retroviruses.

SPRE-like elements were detected in non-human *syncytin* genes; *mac-syncytin-3*, *syncytin-Ten1*, and *syncytin-Car1* (Fig 4A). These SPRE-harboring *syncytin* genes did not form a single clade in a phylogenetic tree of Syncytin and exogenous retroviral Env proteins (Fig 4D), suggesting that the SPRE-like elements emerged in a convergent manner during evolution or were lost in some specific lineage. The more likely scenario is that recombination between retroviruses has led ti the phylogenetic discrepancy between SPRE and Env. SPRE-less *syncytin* genes may also have unrevealed RNA elements. Mouse *syncytin-A* does not require it’s 3′ UTR (Fig 4C) for its expression, whereas it is reported that a ruminant ERV-derived *env* gene, *fematrin-1* does require it’s 3′ UTR for efficient expression (32). Considering the diversity of RNA elements in exogenous retroviruses, it is likely that various RNA elements have been conserved for the efficient expression of ERV-derived genes.

Codon-optimized *syncytin-2* caused cell fusion in the mutant SPRE as much as in the wild-type SPRE. suggesting that the expression of wile-type *syncytin-2* is restricted in a codon usage-dependent manner, and SPRE counteracts the repressive regulation (Fig 6). Similarly, it is known that HIV genes, *gag*, *pol*, and *env* have different codon usages from highly expressing host genes, and importantly, codon-optimization makes their expression Rev/RRE-independent (39, 40). We hypothesized that SPRE-harboring *syncytin* genes have different codon frequencies from other *syncytin* genes. To test this hypothesis, *syncytin* genes were classified according to the relative codon frequency, but no specific relations were observed between SPRE-dependency and codon usages (S3 Fig). It cannot be ruled out the possibility that some repressive RNA motifs in *syncytin-2* ORF were removed by codon-optimization. In HIV-1 *gag* and *pol* gene, multiple negatively acting sequences were detected within un-spliced and partially spliced mRNAs by serial clustered mutagenesis (41). However, it would be difficult to distinguish the effects of the codon frequency from the repressive sequences in the ORF. In either case, these findings revealed the convergent evolution of complex viral RNA regulation between current and ancient retroviruses, in which the repressive regulation intrinsic to viral ORFs are counteracted by other RNA regulatory systems.

The SPRE mutation increased the length of poly(A) tails in *syncytin-1* mRNA (Fig 6). Though the relationship between the poly(A) tail sizes and the translation efficiency has not been observed or only observed in limited developmental stages in early comprehensive studies on the poly(A) tail length (42, 43), a more recent study revealed that short poly(A) size is a feature of highly translated genes (36, reviewed in 44). Though this phenomenon was not observed in *syncytin-2* mRNA, the longer poly(A) tail of *syncytin-1* mRNA would be one of the mechanisms underlying it’s reduced translation efficiency.

The discovery of SPRE implies that ERVs harbor more hidden RNA elements. The co-option of retroviral regulatory elements is also important for the regulation of host genes. LTRs of ERVs contain various *cis*-regulatory elements, enhancers, promoters, and polyadenylation signals, that influence host gene expression, contributing to the diversification of species-specific gene expression in immune systems (45) and placental development (46, 47). By analogy, it is also possible that RNA elements in ERVs and other retrotransposons contribute to the host gene regulatory networks. 27.7% of mouse and 28.5% of human RefSeq transcripts contain at least one retrotransposon in their 3′UTR, and percentages of ERVs and retrotransposons in 3′ UTRs negatively correlate with gene expression levels (48). Though molecular mechanisms of this phenomena are still unclear, the finding of SPRE suggested that ERV-derived elements in 3′ UTRs may affect host gene expression via viral RNA motifs. Some ERV-derived RNAs also function as long non-coding RNAs (lncRNAs). Recent studies have reported that ERV-derived lncRNAs can affect the host gene regulatory networks by binding to host RNA binding proteins (49, 50, 51). Such lncRNAs may harbor RNA elements that have evolved to provide advantages to ancient retroviruses. Therefore, in addition to characterizing transcripts of ERV-derived genes such as *syncytin* genes, the functional analysis of ERV-derived lncRNAs will also provide the opportunity to discover more hidden RNA elements and the new implication to the deep evolution of retroviral RNA elements.

### Materials and Methods Cell cultures

293T (#RCB2202, Riken BioResource Research Center), COS-7 (#CRL-1651, ATCC), NIH3T3-3-4 (#RCB1862, Riken BioResource Research Center), MDCK (#CCL-34, ATCC), QT6 (#CRL-1708, ATCC) were cultured in Dulbecco’s modified Eagle’s medium (#D5796, Sigma Aldrich) supplemented with 10% heat-inactivated fetal calf serum (#10270106, Gibco), and penicillin (100 units/mL) and streptomycin (100 mg/mL) (#09367-34, nacalai tesque) at 37°C in a humidified atmosphere of 5% CO_2_ in air.

### Plasmids

The Syncytin-1 and Syncytin-2 expression plasmids of phCMV3 backbone (#P003300, Genlantis) were constructed previously (20). For minigene construction, ERVWE1 and ERVFRDE1 loci were inserted into the SmaI site of pSP73 (#P2221, Promega) with the fragments of cytomegalovirus (CMV) enhancer and promoter amplified from pcDNA3.1 (#V79020, Thermo Fishier Scientific) using NEBuilder HiFi DNA Assembly Master Mix (#M5520AA, NEB). For the site-directed mutagenesis, linearized vectors with mutations were generated by inverse PCR and they were joined and cyclized by NEBuilder. For the construction of HiBiT reporter plasmids, HIV-1 Gag from pHG (13) was amplified by PCR and inserted into the EcoRI and BamHI sites of phCMV3, and the HiBiT sequence was inserted into the C-terminus by inverse PCR followed by NEBuilder. HIV-1 *gag* was replaced to the MLV *gag* amplified from pGag-Pol-IRES-bsr (52) and the SFV *gag* amplified from pJM356 (53). For the NanoLuc expression plasmid, a fragment of NanoLuc gene were amplified from pNL1.1 (#N1001, Promega) and inserted into the HindIII and EcoRI sites of phCMV3. SPRE-like elements were synthesized and inserted between the EcoRI and NotI sites of each HiBiT reporter plasmid. For the construction of codon-optimized *syncytin-2* minigenes, dsDNA coding codon-optimized *syncytin-2* was synthesized by Eurofins Genomics (Tokyo, Japan). Synthesized DNA was amplified and replaced to the ORF of *syncytin-2* in pFRDE1. All PCRs described above were carried out using KOD One Master Mix (#KMM-101, TOYOBO) with a C1000 Touch thermal cycler (Bio-Rad). Primer sequences are listed in S3 Table, and representative plasmid sequences are included in S1 Text.

### Fusion-dependent luciferase assay

A fusion-dependent luciferase assay was performed as described previously (54) with some modifications. 293T cells were seeded onto 24-well plates (2.5 × 10^5^ cells/well). On the next day, cells were co-transfected with each Syncytin expression plasmid (500 ng), pT7EMCLuc (500 ng), and pRL-TK (50 ng) with 0.7 µl of Avalanche Everyday Transfection Reagent (#EZT-EVDY-1, EZ Biosystems) according to the manufacturer’s instruction. Six hours post-transfection, cells were resuspended and cocultured with 293T cells transfected with a T7 polymerase expression vector, pCAGT7. Then, 24 hours after coculture, luciferase activities of cell lysates were measured using the Dual-Luciferase Reporter Assay System (#E1910, Promega). All experiments were conducted more than three times independently.

### HiBiT luciferase assay

293T, COS-7, NIH3T3-3-4, MDCK, and QT6 cells seeded in 24-well plates (1.2-2.5 × 10^5^ cells/well) were transfected with HiBiT reporter plasmid (500 ng) with Avalanche Everyday Transfection Reagent. Twenty-four hours after transfection, cells were subjected to luciferase assay using Nano Glo HiBiT Lytic Detection System (#N3030, Promega).

### Northern blot analysis

For sample preparation, two µg of each minigene plasmid was transfected to 293T cells seeded in 6-well plates (1 × 10^6^ cells/well). Twenty-four hours after transfection, total RNA was extracted using RNAzol RT (#RN109, Molecular Research Center) and stored at −80°C. Detailed protocol and recipe of the reagents for Northern blotting were followed by the manufacture’s protocol, “DIG application manual for Filter Hybridization” (Roche). Briefly, for probe construction, DNA fragments with T7 promoter were obtained by PCR with T7 promoter sequence-attached primers listed in S3 Table. The amplicons were used for templates of *in vitro* transcription using DIG RNA Labeling Kit (#11175025910, Roche). RNA samples were mixed with 2 × Loading Buffer and heated at 65°C for 10 min. Then, Each RNA sample (2.5 µg/lane) was loaded into 1.5% agarose gel in 1 × morpholine propane sulfonic acid buffer containing 2% formaldehyde. After electrophoresis with 100 V for 180 min, samples were transferred onto the positively charged nylon membrane (#11209272001, Roche) overnight and fixed by UV cross-linking. DIG-labeled RNA probes were hybridized with DIG Easy Hyb (#11796895001, Roche) at 68°C overnight, and anti-Digoxigenin-AP, Fab fragments from sheep (#11093274910, Roche) were bound to probes with Blocking Reagent (#11096176001, Roche). Signal detection was performed by CDP-Star (#11685627001, Roche) with LAS-4000 (FUJIFILM).

### Poly(A) tail-targeted PCR

Total RNA was isolated using RNAzol RT from 293T cells 24 hours after transfection. Universal miRNA Cloning Linker (#S1315S, NEB) was linked to the 3′-terminus of RNA using T4 RNA Ligase 2, truncated K227Q (#M0351S, NEB). Then, cDNA was synthesized using Verso cDNA Synthesis Kit (#AB1453, Thermo Fisher Scientific) and PCR was performed using KOD One PCR Master Mix (#KMM-101, TOYOBO). Resulting PCR fragments were cloned into pSP73 and verified by sanger sequencing. Primers are listed in S3 Table.

### Sequence search for SPRE

Nucleotide sequences of 6,915 repeat elements were retrieved from Dfam 3.2 curated families (July/2020) (https://www.dfam.org/home) (22). For the first step search, the 17-nt SPRE-core motif (5′-TCAGCAGGAAGCAGTTA-3′) was searched by SSEARCH version 36.3.8 in FASTA3 programs (55) in Dfam sequences classified as ERV1, ERV2, and ERV3. To calculate the relative position score, each nucleotide position of the hit to the SPRE-core motif was normalized by the nucleotide length of each ERV sequence. The hits to the 17-nt core motif which were located in ERVs belonging to ERV1 group and whose relative position scores were more than 97 were collected with their 5′ flanking (30-nt) and 3′ flanking (42-nt) sequences. Then, they were aligned using MAFFT version b7.402 (56). The alignment was subjected to HMM-based search in the 6,915 Dfam entries using nhmmer in HMMER vsertion3.3.1 (23) for the second step search. The alignment of the resultant SPRE-like elements was output using nhmmer with -A option. The alignment was visualized by AliView version1.23 (57) and WebLogo3 (58). To search for the SPRE-like elements in host genomes, genomic sequences of 318 species of mammals and 499 species of birds available in the NCBI genome database as of December 13^th^, 2020 (https://www.ncbi.nlm.nih.gov/genome/) were retrieved. To search for the SPRE-like element in exogenous viruses, all viral sequences in NCBI nucleotide collection including partial viral sequences were retrieved from NCBI virus with taxonomy ID:10239 (https://www.ncbi.nlm.nih.gov/labs/virus/) at October 13^th^, 2020. Nhmmer was used for the above sequence search with the HMM profile constructed from the alignment of 41 SPRE-like elements. A phylogenetic tree of taxa was constructed by TimeTree (http://www.timetree.org/) (59).

### Construction of a phylogenetic tree

MEGA X software suite (60) was used for our phylogenetic analyses as follows. Since the transmembrane (TM) domain of Env proteins is known to be more conserved than the surface (SU) domain in many retroviruses, the amino acid sequences of the TM domain were used for the phylogenetic analysis. The amino acid sequences were aligned by MUSCLE program with default option. Maximum likelihood tree was generated with WAG+F model as suggested by the MEGA X model selection. The robustness of the phylogenetic tree was evaluated by 1000 bootstrap duplicate data sets.

### Analysis of codon usages

Relative codon frequency of *syncytin* genes and reporter genes were calculated by Codon Usage Generator v2.4 (61, 62). Heatmap was generated by heatmap function of R version 4.02.

## Data Availability

All relevant data are within the manuscript and its Supporting Information files.

## Author contributions

KK and TM designed research; KK performed experiments; KK and SN analyzed data; and KK, SN, and TM wrote the paper.

## Acknowledgement

We are grateful to Rachael Tarlinton (University of Nottingham) for helpful advice to the manuscript.

---

**S1 Fig.**
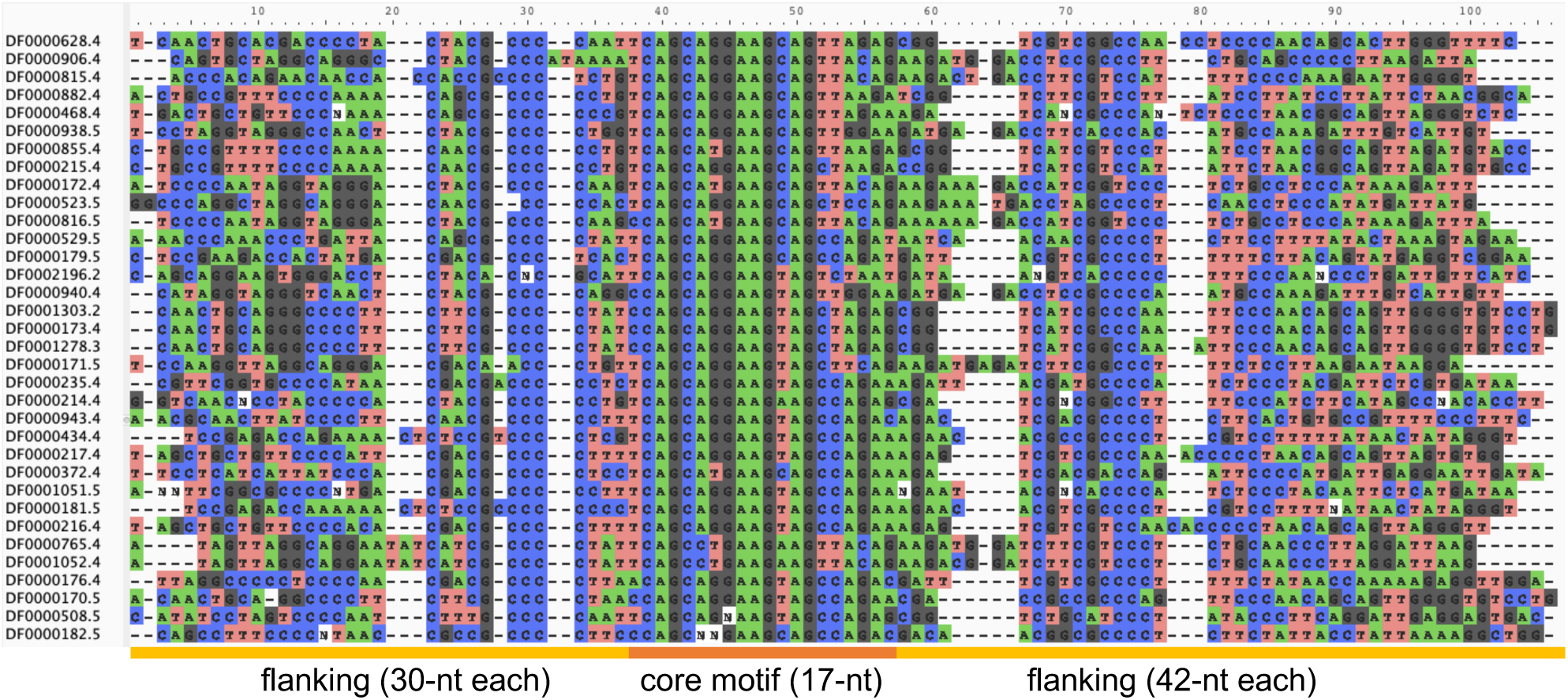
Alignment of the SPRE-like elements detected in the first step search (see Fig 2A).

**S2 Fig.**
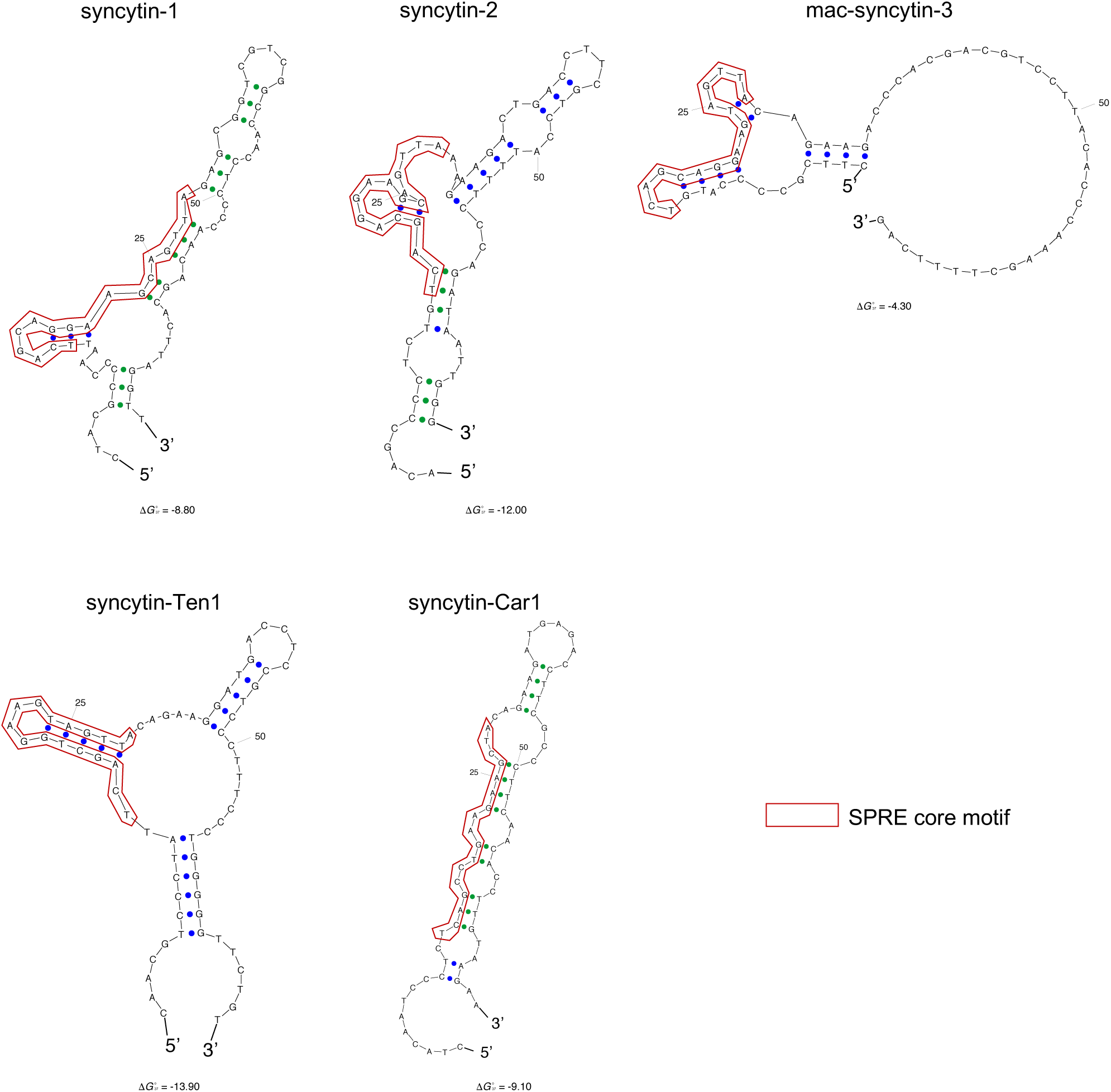
RNA secondary structures of the SPRE-like elements. Structure predictions were conducted by Sfold webserver (http://sfold.wadsworth.org/cgi-bin/srna.pl) with default parameters.

**S3 Fig.**
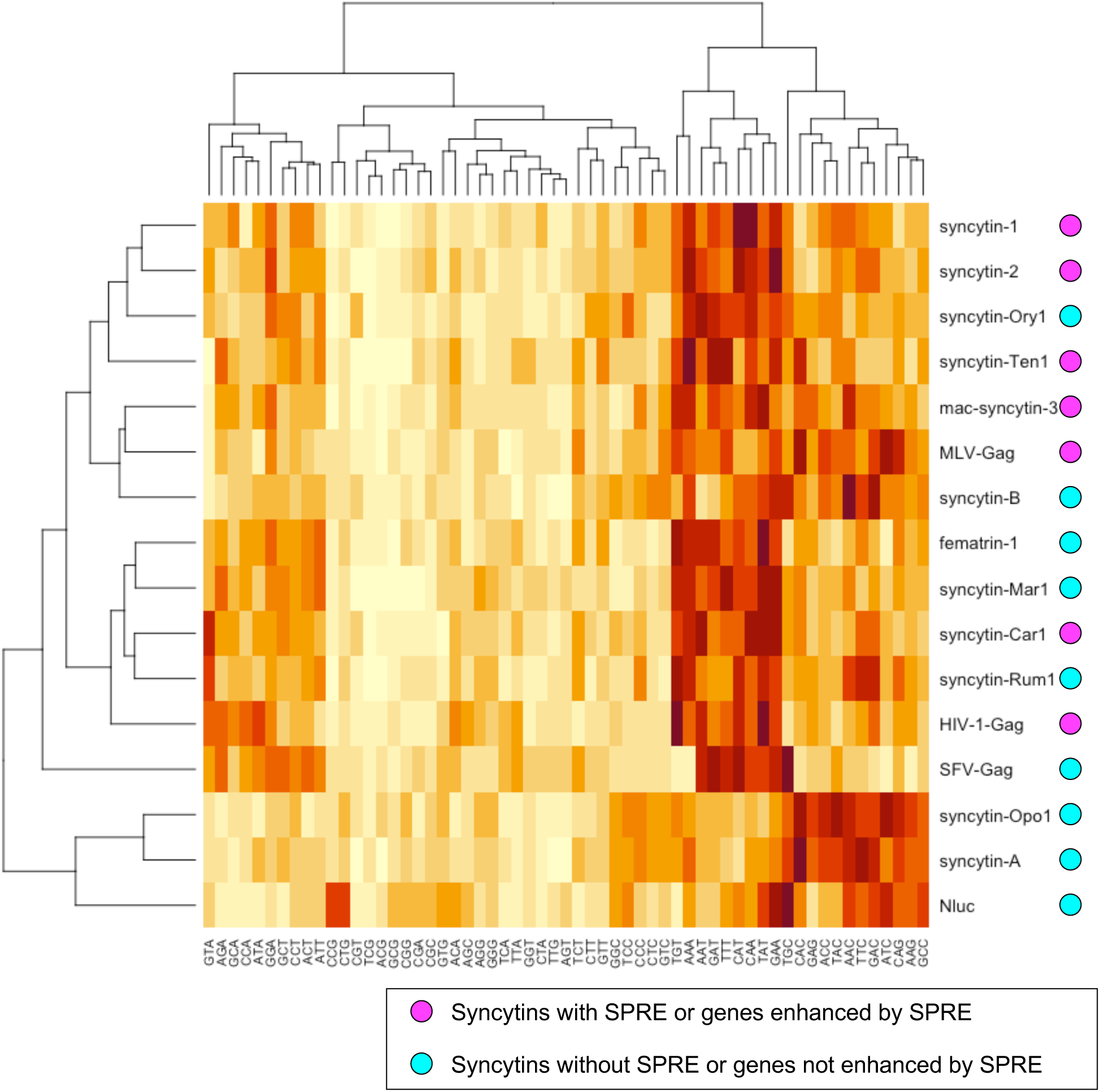
Heatmap classification of *syncytin* and reporter genes by their codon frequencies.

**S1 Table. Results of SPRE search in Dfam database by SSEARCH. (XLSX)**

**S2 Table. Results of SPRE search in mammalian and avian genomes. (XLSX)**

**S3 Table. Lists of primers and accession numbers. (XLSX)**

**S1 Text. Representative plasmids sequences. (TXT)**

